# Synchronous spiking associated with high gamma oscillations in prefrontal cortex exerts top-down control over a 5Hz-rhythmic modulation of spiking in locus coeruleus

**DOI:** 10.1101/2020.04.26.061085

**Authors:** Nelson K. Totah, Nikos K. Logothetis, Oxana Eschenko

**Author notes:** Corresponding author: Nelson K. Totah or Oxana Eschenko. Author Contributions: Conceptualization – NKL, NKT, OE Methodology – NKT Software – NKT Formal Analysis – NKT Investigation – NKT Resources – NKL Writing: Original Draft – NKT Writing: Review and Editing – NKT, OE, NKL Visualization – NKT Supervision – OE Funding Acquisition – NKL.

## Abstract

The brainstem noradrenergic locus coeruleus (LC) is reciprocally connected with the prefrontal cortex (PFC). Strong coupling between LC spiking and depolarizing phase of slow (1 – 2 Hz) waves in the PFC field potentials during sleep and anesthesia suggests that the LC drives cortical state transition. Reciprocal LC-PFC connectivity should also allow interactions in the opposing (top-down) direction, but prior work has only studied prefrontal control over LC activity using direct electrical (or optogenetic) stimulation paradigms. Here, we describe the physiological characteristics of naturally occurring top-down prefrontal-coerulear interactions. Specifically, we recorded LC multi-unit activity (MUA) simultaneously with PFC single unit and local field potential (LFP) activity in urethane-anesthetized rats. We observed cross-regional coupling between the phase of ~5 Hz oscillations in LC population spike rate and the power of PFC LFP oscillations within the high Gamma (hGamma) range (60 – 200 Hz). Specifically, transient increases in PFC hGamma power preceded peaks in the ~5 Hz LC-MUA oscillation. Analysis of cross-regional transfer entropy demonstrated that the PFC hGamma transients were predictive of a transient increase in LC-MUA. A ~29 msec delay between these signals was consistent with the conduction velocity from the PFC to the LC. Finally, we showed that PFC hGamma transients are associated with synchronized spiking of a subset (27%) of PFC single units. Our data suggest that, PFC hGamma transients may indicate the timing of the top-down excitatory input to LC, at least under conditions when LC neuronal population activity fluctuates rhythmically at ~5 Hz. Synchronized PFC neuronal spiking that occurs during hGamma transients may provide a previously unknown mode of top-down control over the LC.

## Introduction

A common assumption about coerulear-prefrontal (LC-PFC) functional connectivity is that the LC is a driver. This assumption is based on the well-documented actions of the LC as an ascending neuromodulatory system (Swanson and Hartman, 1975; Grzanna et al., 1977; Fallon et al., 1978; Morrison et al., 1979; Loughlin et al., 1982; Waterhouse et al., 1983; Schwarz et al., 2015; Kebschull et al., 2016). However, bidirectional LC-PFC interaction is also likely as the LC and PFC are reciprocally and monosynaptically connected. Indeed, the PFC has been demonstrated to exert both inhibitory and excitatory effects on LC activity (Arnsten and Goldman-Rakic, 1984; Sesack et al., 1989; Luppi et al., 1995; Sara and Hervé-Minvielle, 1995; Jodo et al., 1998; Aston-Jones and Cohen, 2005; Breton-Provencher and Sur, 2019). Notably, the PFC is the only cortical region sending direct projections to LC (Sesack et al., 1989; Luppi et al., 1995). Previous studies on LC-PFC interactions during sleep or anesthesia have focused on a prominent 1 - 2 Hz oscillation in LC spike rate that is thought to promote the transition to cortical heightened excitability (Eschenko et al., 2012; Safaai et al., 2015; Totah et al., 2018). However, during urethane anesthesia, rhythmic LC activity occurs not only at ~1 - 2 Hz, but also at ~5 Hz (Safaai et al., 2015). Here, we studied the nature and the directionality of the LC-PFC interaction during these faster ~5 Hz fluctuations of LC multi-unit activity (MUA).

In the present study, we monitored LC-MUA and wide-band extracellular activity from the prelimbic division of the prefrontal cortex (PFC) in urethane-anesthetized rats. Importantly, while recording LC-MUA for long durations and with stable spiking activity in behaving animals continues to present an immense challenge, anesthesia permits stable, long-lasting recordings to study physiological interactions between the LC and PFC. Here, we report cross-regional phase-amplitude coupling between LC-MUA ~5 Hz oscillations and high gamma (hGamma, 60-200 Hz) LFP power in the PFC. Transient increases in PFC hGamma power preceded LC-MUA ~5 Hz oscillation peaks. hGamma transients were associated with PFC unit-pair spike synchrony. Taken together, our results demonstrate that, during epochs when LC population firing rate oscillates at ~5 Hz hGamma transients are a sign of PFC top-down excitatory control over the LC.

## Results

Our goal was to study the nature and directionality of LC-PFC interactions during epochs when LC population firing rate oscillated at ~5 Hz. For this purpose, we used urethane-anesthetized rats, a common model for studying LC-PFC interactions (Sara and Hervé-Minvielle, 1995; Jodo et al., 1998; Marzo et al., 2014; Safaai et al., 2015; Neves et al., 2017; Totah et al., 2018). We recorded wide-band (0.1 Hz – 8kHz) extracellular activity from deep layers of the prelimbic division of the rat PFC and from the LC core. LC-MUA was measured by first band-pass filtering (400 – 3 kHz) to resolve extracellular spiking and then rectifying the signal. **Figure 1A** shows an example trace of band-pass filtered extracellular spiking signal (grey line) and the rectified LC-MUA signal (purple line). Large amplitude fluctuations in LC-MUA are generated primarily by action potentials produced by the neuronal population within 300 um of the electrode (Logothetis, 2003, 2008). This recording radius is comparable with the smallest dimension of LC core (Grzanna and Molliver, 1980). Therefore, MUA was likely only capturing LC neuronal activity. We verified that the MUA originated from LC norephinephrine (NE) containing neurons by injecting clonidine (0.05 mg/kg, i.p.) at the end of each recording session. Clonidine completely abolished the extracellular spiking that contributes to MUA signal in all rats (an example rat is shown in **Figure 1B**); therefore, clonidine administration demonstrated that the LC-MUA signal was confined to the LC nucleus and consisted of spikes emitted by the LC-NE neurons.

**Figure 1.**
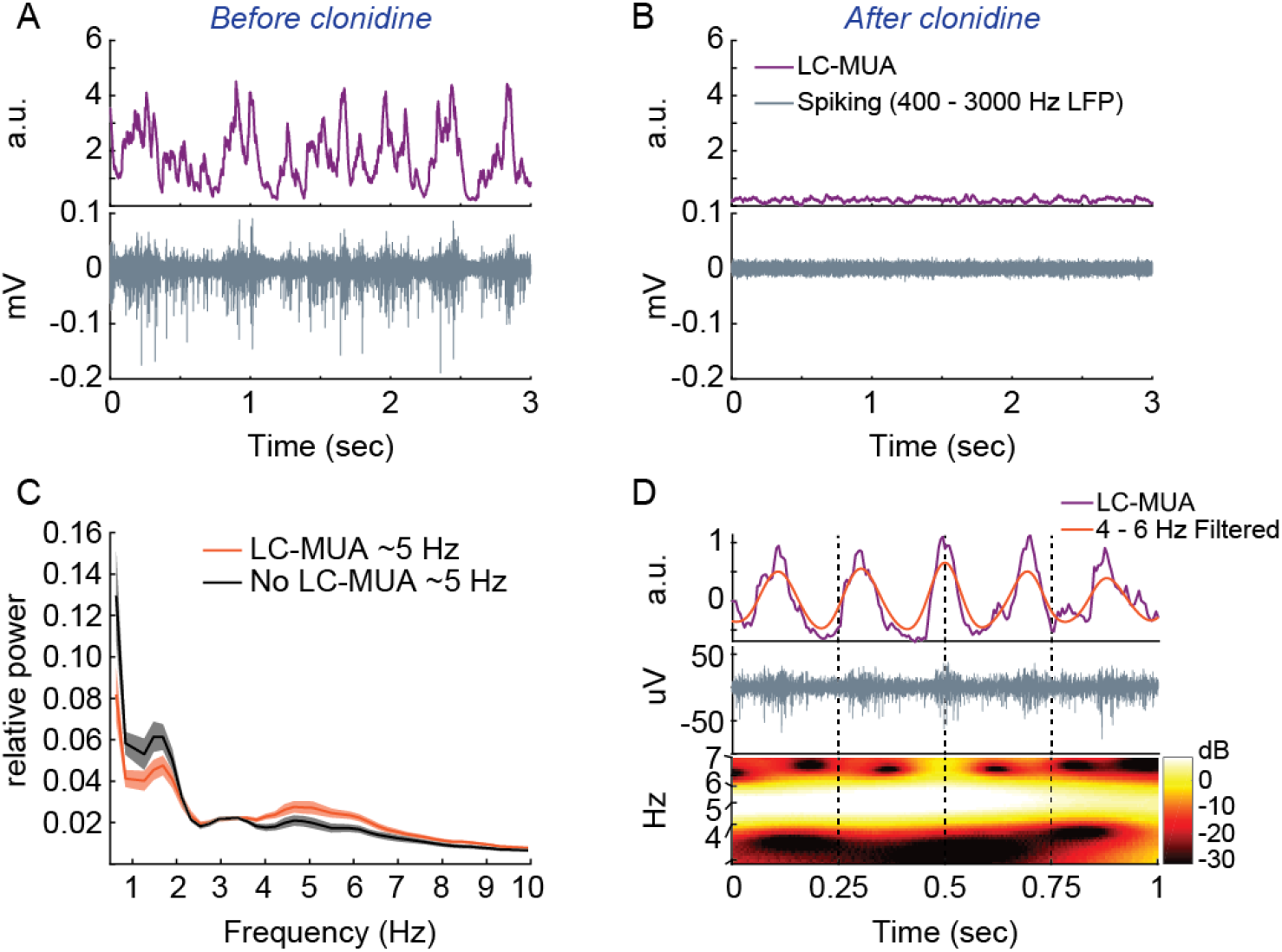
Multi-unit activity (MUA) in the LC exhibited rhythmic 5 Hz fluctuations. **(A)** High-pass filtered (> 400 Hz) extracellular activity (grey line) recorded from the LC. The band limited power (purple line) was obtained by rectifying the 400 – 3000 Hz bandpass filtered signal. **(B)** Systemic administration of clonidine caused cessation of LC-MUA. Clonidine inhibits LC norepinephrine (NE) neurons by binding to alpha-2 auto-inhibitory adrenoreceptors present on the soma and dendrites of LC-NE neurons (Aghajanian et al., 1977). Clonidine administration discriminates extracellular unit spiking by LC-NE neurons from surrounding non-LC neurons because structures in the vicinity of the recording electrode do not have alpha-2 receptors (McCune et al., 1993). **(C)** The average LC-MUA power spectrum (normalized by total power) during epochs with and without LC-MUA ~5 Hz oscillations (N = 25 out of 35 rats). Each 4 sec recording epoch was classified as LC-MUA 5 Hz or non-5 Hz epoch and averaged within rat. The plots present the grand average across rats with standard error shown as shading. **(D)** An example of LC-MUA 5 Hz oscillatory activity. The grey line is the high-pass filtered LC-MUA (>400 Hz). The purple line is the band limited power (purple line) of the 400 - 3000 Hz bandpass filtered signal, as in panels A and B. The orange line is the 4 – 6 Hz filtered LC-MUA. The wavelet transform of the purple line (LC-MUA) shows a clear 4 – 6 Hz oscillation.

### The frequency-specific modulation of the PFC activity during LC population oscillations at ~5Hz

Consistent with an earlier report (Safaai et al., 2015), we confirmed that LC-MUA oscillates at both ~1 - 2 Hz and ~5 Hz during urethane anesthesia. We characterized LC-MUA oscillations by calculating the power spectral density (PSD) of the LC-MUA. For each recording session (n = 35 rats), we calculated the PSD in 4-sec epochs and clustered them using principle components analysis and k-means clustering. Epochs with ~5 Hz oscillations of LC-MUA were identified as a cluster with a peak in the 4 - 7 Hz range. We detected these epochs in 25 out of 35 recording sessions (n = 25 rats). **Figure 1C** shows the average power spectrum of all 4-sec data epochs with LC-MUA 5 Hz oscillations versus epochs without LC-MUA 5 Hz oscillations. **Figure 1D** shows an example clip of LC population rhythmic firing at ~5 Hz.

Prior research has demonstrated that slow (1 - 2 Hz) rhythmic LC activity occurs during sleep and anesthesia when the cortex is in a slow oscillation state in the same frequency range (Sara and Hervé-Minvielle, 1995; Lestienne et al., 1997; Eschenko et al., 2012; Safaai et al., 2015; Totah et al., 2018). However, the brain state during which LC-MUA 5 Hz oscillations occur is not known. We assigned each 4-sec recording epoch with LC-MUA 5 Hz oscillations to a “slow oscillation” or an “activated” cortical state (see Methods for cortical state classification). The slow oscillation state consisted of periodically (1 - 2 Hz) alternating epochs of high and low neuronal excitability, whereas the “activated” state was one of continuously enhanced neuronal excitability (**Figure 2A**). LC-MUA 5 Hz oscillations occurred mostly during the cortical activated state (**Figure 2B**). Significantly more epochs of LC-MUA 5Hz were observed during the cortical activated state in comparison to the slow oscillation state (χ^2^ = 3494.7, p<0.0001). Having observed a brain state-dependency of LC-MUA 5 Hz oscillations, we focused the remaining analyses on the epochs that occurred during the activated cortical state.

**Figure 2.**
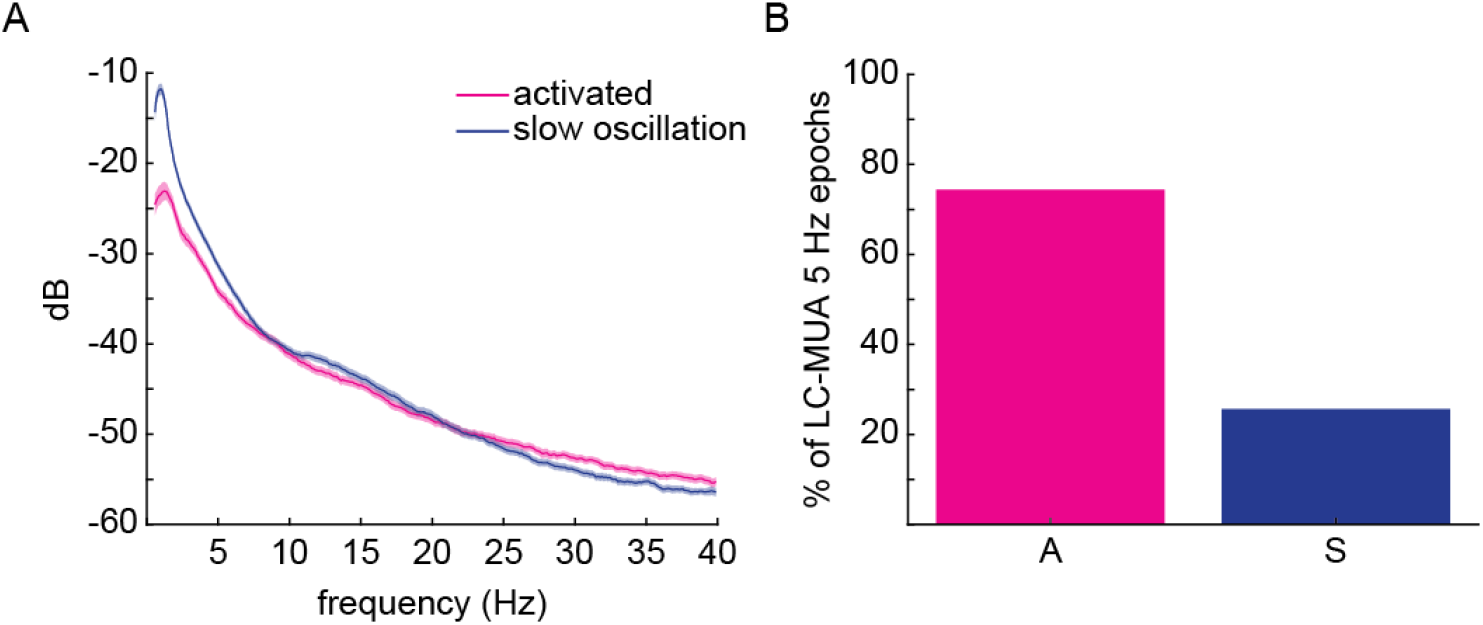
LC-MUA 5 Hz oscillations occurred primarily in the activated cortical state. **(A)** The PFC LFP power spectrum for the activated and slow oscillation states. The lines are the means across rats and the shading is the standard error. **(B)** The percentage of LC-MUA 5 Hz epochs occurring in each cortical state.

Although a phasic increase in LC-MUA has been proposed as a driver of the cortical activated state, the nature and directionality of LC-PFC interactions during epochs when LC population activity oscillates at ~5Hz has not yet been characterized. We first measured the relationship between the phase of LC-MUA 5 Hz oscillations (i.e., relative increases and decreases in LC population spike rate) and changes in the power spectrum of the PFC LFP. This relationship was quantified using a modulation index (MI) that measured the non-uniformity of the phase distribution of PFC LFP amplitude between 30 and 300 Hz (Tort et al., 2010). Given that the PFC projections to the LC originate in the deeper (3/5) layers and the likelihood that this PFC neuronal population influences LC activity, we targeted the PFC deep layers (Sesack et al., 1989; Luppi et al., 1995). The analysis revealed that LC-MUA 5 Hz oscillations are associated with frequency band-specific modulation of PFC LFP power between 60 Hz and 200 Hz. This band includes high gamma (hGamma) as well as high frequency oscillations (HFOs) (Ray et al., 2008a, Ray et al., 2008b; Ray et al., 2011; Khodagholy et al., 2017). We will refer to this range (60 – 200 Hz) as the hGamma band, although it also includes HFOs. **Figure 3A** shows the average MI value across all recording sessions in which LC-MUA 5 Hz oscillation epochs were present during the cortical activated state. Nine rats with less than 10 epochs (total duration 40 sec) in the activated state were excluded, leaving 16 rats. We excluded further 4 sessions (n = 4 rats) due to a lack of clear modulation in the PFC power spectrum that was inconsistent with the population mean shown in **Figure 3A**; thus, 12 rats were assessed. **Supplementary Figure 1** illustrates the MI plots for the 4 excluded sessions as well as examples of 4 accepted sessions for comparison. The mean MI value for all 4 - 7 Hz epochs was significant (i.e., MI>0) in all 12 accepted sessions (n = 12 rats) (**Figure 3A**). A boxplot illustrates the distribution, across rats, of the average MI value for 4 – 7 Hz phase with hGamma (60 – 200 Hz) amplitude (**Figure 3B**). The temporal relationships between the PFC hGamma amplitude and LC-MUA 5 Hz oscillation phase are shown on PFC LFP power spectrograms triggered on the peaks of the LC-MUA 5 Hz oscillation (**Figure 3C**). Consistent temporal relations between the LC-MUA rhythmic increases at 5 Hz and PFC LFP power increases exclusively in the hGamma band contrasts with prior work demonstrating LC activation triggering a less specific (>30 Hz) change in PFC LFP (Marzo et al., 2014; Neves et al., 2017).

**Figure 3.**
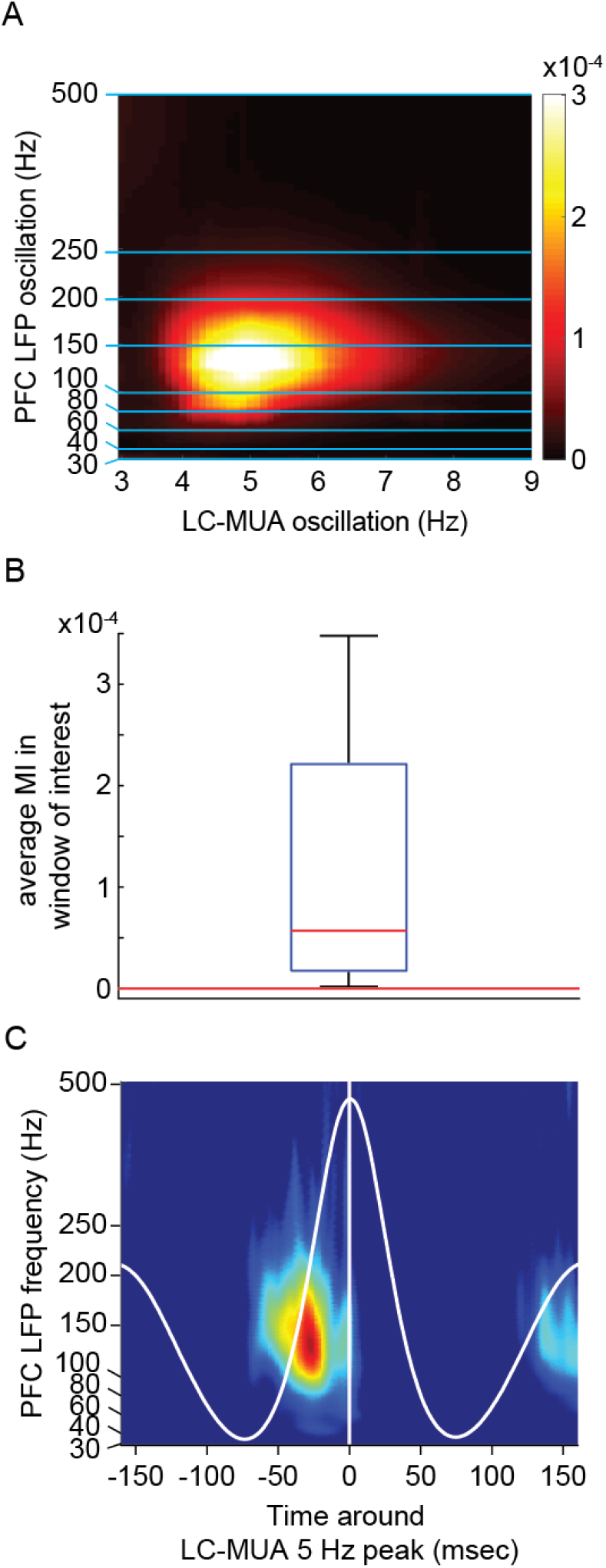
The phase of LC-MUA 5 Hz oscillations was associated with a frequency-specific (60-200Hz) modulation of PFC LFP. **(A)** The average modulation index (MI) is plotted for each LC-MUA oscillation phase against PFC LFP oscillation amplitude (N = 12 rats). Zero values (black) are not significantly higher than those expected by chance (one-sided permutation test, p<0.01). **(B)** The box plot shows the distribution of Mis, across rats, in the window of interest (4 to 7 Hz phase, 60 to 200 Hz amplitude). For each rat, the values in this window of interest were not normally distributed (Shapiro-Wilk test) and the mean was influenced by very high values (i.e., the “hot spot” in panel A). Since we wanted to quantify this the magnitude of this hot spot across rats, we used the mean, rather than the median, to obtain a summary MI value for each rat. The box plot shows the distribution of the MI hot spot magnitude across rats. Two outlier MIs (highly significant), which were 9.5E-4 and 5.4E-4, are not shown on the box plot to allow visualization of the distribution of data. Note that MI was greater than 0, which indicates significant cross-frequency coupling was obtained for all 12 accepted sessions (N = 12 rats). **(C)** The PFC LFP power spectrogram is plotted aligned to the peak of LC-MUA 5 Hz oscillation. The spectrogram was first averaged across LC-MUA 5 Hz peaks and then averaged across rats. The white line shows LC-MUA (4 – 6 Hz filtered) aligned to peaks and averaged over all accepted sessions. The PFC LFP power is Z-score normalized within each frequency bin. The hGamma power increase preceding the peak of LC-MUA 5 Hz oscillation is apparent.

**Supplementary Figure 1.**
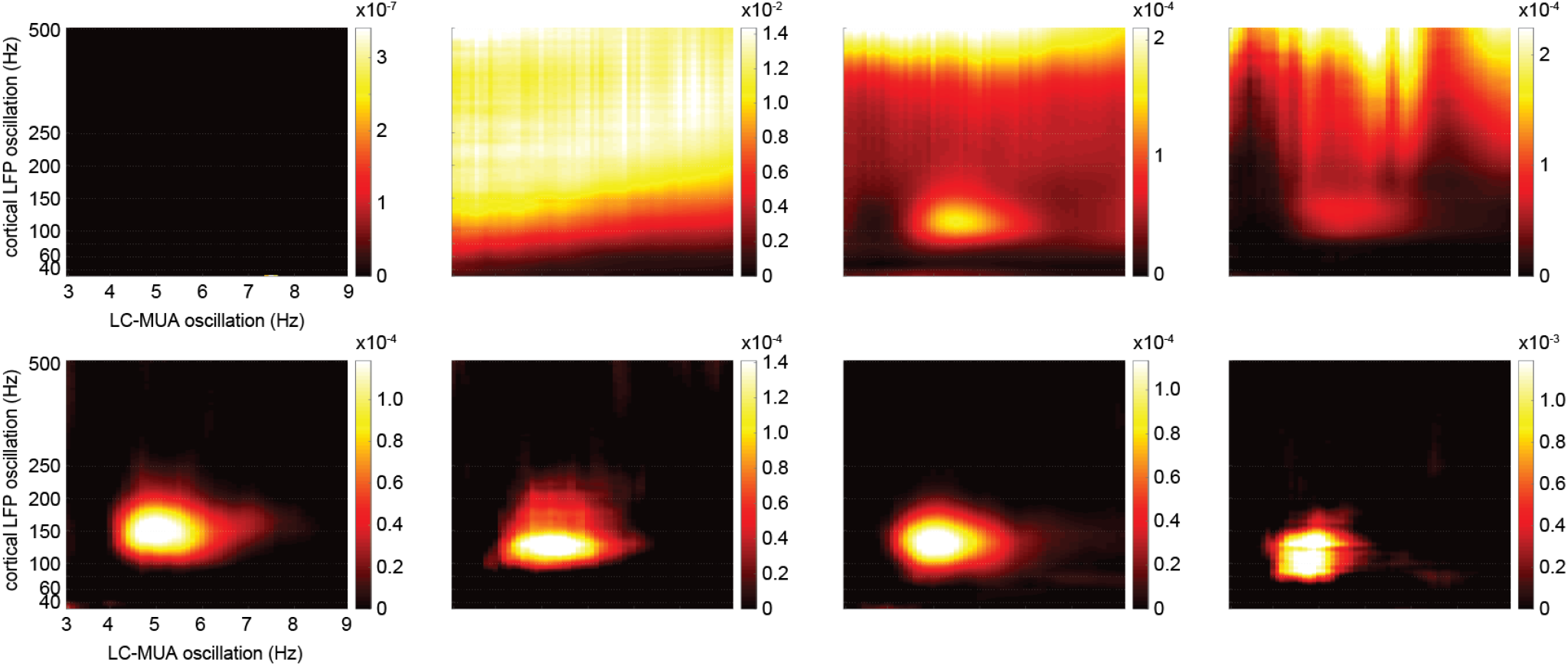
The MI is shown for individual example sessions. The top row shows 4 excluded sessions due to the pattern deviating from accepted cases (4 example shown on the bottom row).

### The directionality of the LC-PFC interaction

Having established that transient increases in PFC hGamma power are phase-locked to LC-MUA 5 Hz oscillations, we turned to assessing the directionality of this interaction. It is commonly assumed that the LC, as an ascending neuromodulatory system drives changes in the cortex (Swanson and Hartman, 1975; Grzanna et al., 1977; Fallon et al., 1978; Morrison et al., 1979; Loughlin et al., 1982; Waterhouse et al., 1983; Schwarz et al., 2015; Kebschull et al., 2016, Marzo et al., 2014; Neves et al., 2017). However, the PFC can also exert both inhibitory and excitatory influences on LC activity (Sara and Hervé-Minvielle, 1995; Jodo et al., 1998). In order to characterize the directionality of the LC-PFC interaction during epochs of LC-MUA 5 Hz oscillations, we used information theoretic measures to calculate the transfer entropy (TE) from the phase of the LC-MUA 5 Hz signal to the amplitude of the PFC LFP hGamma signal, as well as PFC to LC (Besserve et al., 2010, 2015). We observed that the direction of interaction during LC-MUA 5 Hz oscillations was actually predominantly from the PFC to the LC (**Figure 4A**). Transfer entropy was larger from the past of the PFC LFP hGamma amplitude to present LC-MUA 5 Hz phase than vice versa (Z = 4.128, p = 3.66E-5, power = 1.0, Cohen’s D = 2.833). This result is consistent with the peri-event spectrograms presented in **Figure 3C**, which show the median latency from the PFC hGamma power modulation to the LC-MUA 5 Hz oscillation peak is 29.1 msec (**Figure 4B**). This delay after the hGamma transient is consistent with the previously reported glutamatergic conduction time from the prelimbic division of the PFC to the LC in the rat (average 35 msec, range 10 to 70 msec (Jodo et al., 1998). Our findings, taken together with the latencies reported in Jodo et al. (1998), suggest that PFC hGamma transients may indicate the timing of the top-down excitatory input to LC.

**Figure 4.**
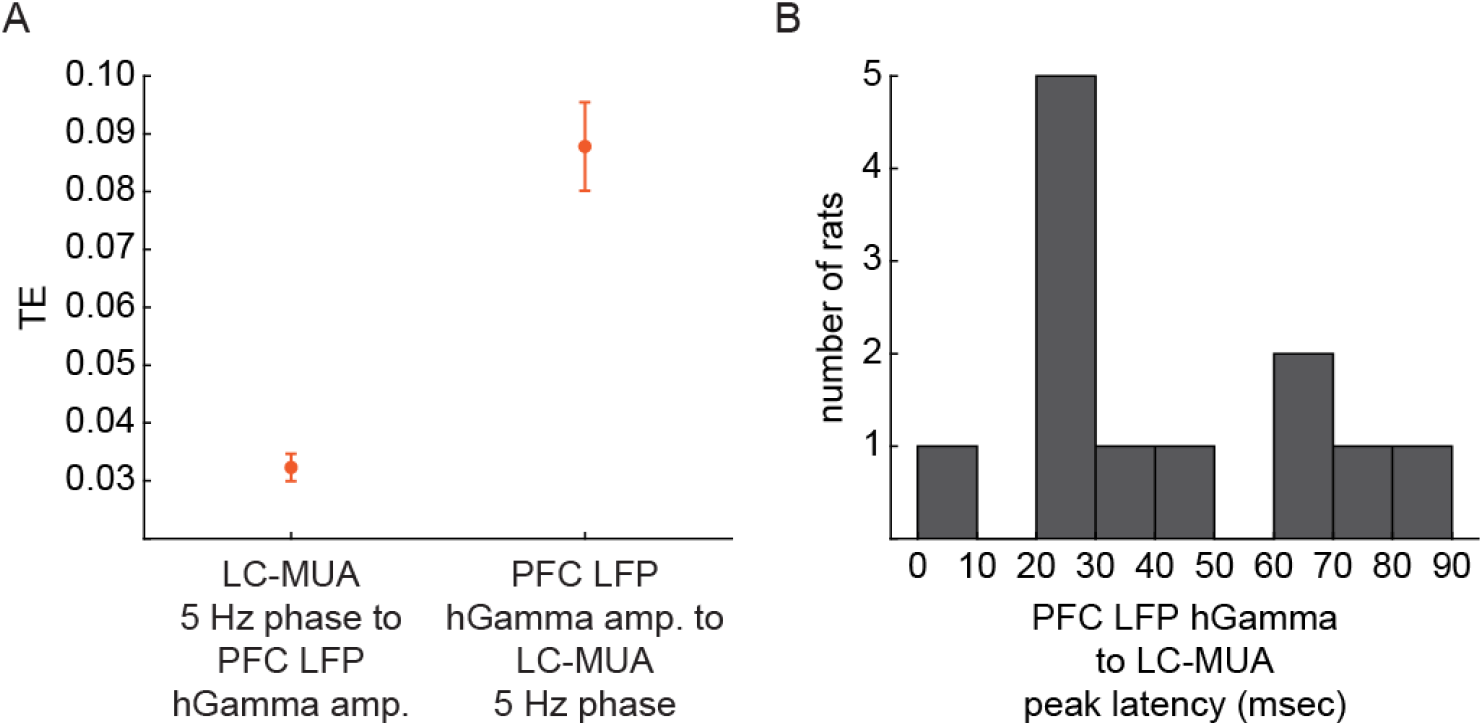
PFC hGamma amplitude is predictive of the future phase of LC-MUA 5 Hz oscillations. **(A)** Transfer entropy (TE) is higher in the direction from PFC hGamma amplitude to LC-MUA 5 Hz phase. The plot shows the mean and standard error of TE for each direction of interaction. **(B)** A histogram showing the latency from the PFC LFP hGamma power peak until the LC-MUA 5 Hz oscillation peak for 12 rats. The median is 29.1 msec with a standard deviation of 25.7 msec and a range of −87.8 msec to −0.9 msec.

The source of the cross-frequency LC-PFC interaction during epochs when the LC population oscillates at ~5 Hz is unknown. However, the LFP signal, which reflects peri-synaptic input around the electrode, did not have a 5 Hz oscillation in the PFC (**Figure 5**).

**Figure 5.**
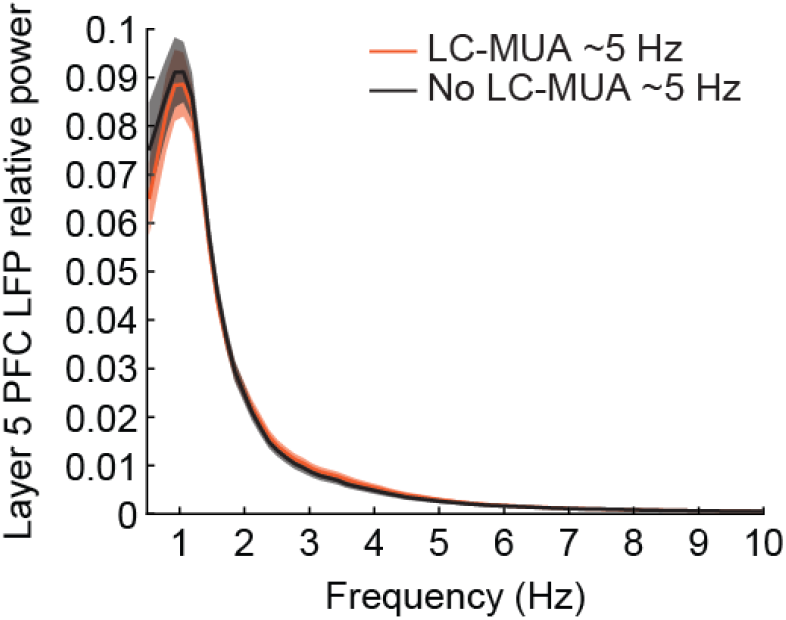
The PFC LFP spectrogram does not contain a peak at ~5 Hz. The power spectrum is plotted separately for epochs with (orange line) and without (black line) LC-MUA 5 Hz oscillations. Shading is the standard error around the mean.

### The PFC spike rate during LC-PFC interactions

The extracellular potential changes recorded in PFC as hGamma oscillations are highly localized and cannot directly affect LC neurons; rather, it is the spiking output of PFC neurons that drives LC activity. We next assessed the spike rate of PFC units during LC-MUA 5 Hz oscillations. In order to assess how PFC spiking and hGamma power relate to 5 Hz rhythmic fluctuations of LC-MUA, we aligned PFC multi-unit spike rate to the peaks of the LC-MUA 5 Hz oscillation. We highpass filtered the wideband PFC extracellular signal (at 500 Hz), detected spike times by thresholding the signal (3.5 standard deviations from the noise), and constructed a SDF from those spike times using a 100 msec Gaussian kernel. We found that PFC MUA was modulated around LC-MUA 5 Hz oscillation peaks, albeit with a slight phase shift compared to PFC hGamma power (**Figure 6A**).

**Figure 6.**
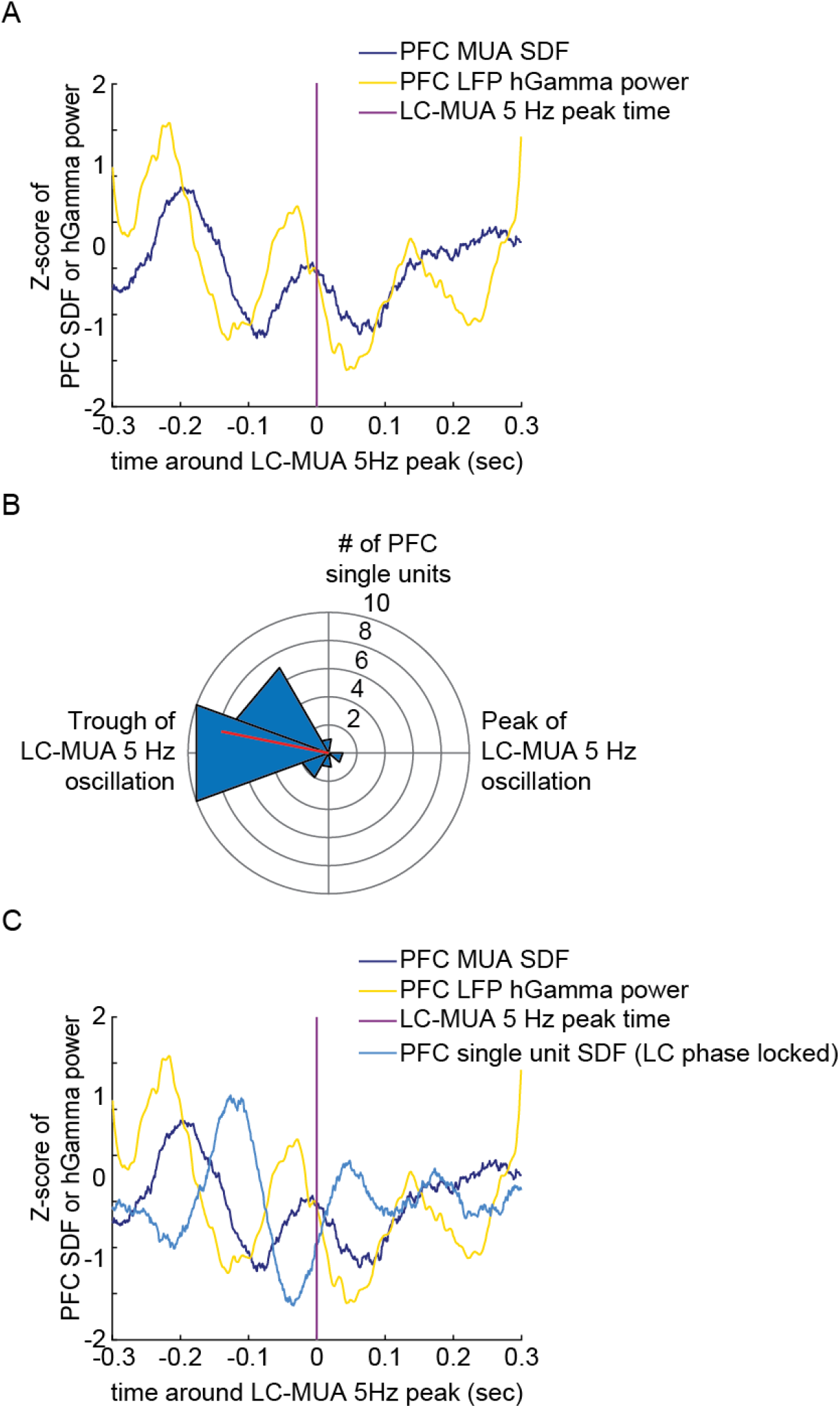
PFC spiking is phase locked to LC-MUA 5 Hz oscillation. **(A)** PFC single unit and multi-unit SDFs and hGamma gamma amplitude co-fluctuate around the LC-MUA 5 Hz peak (purple line). Values have been z-scored to the mean and standard deviation of the entire recording. **(B)** Spike timing of a sub-set of PFC single units (27%) is phase-locked to the tough of the LC-MUA 5 Hz oscillation. The preferred phase is plotted for significantly phase-locked PFC units. The red line shows the circular mean across these phase-locked units. **(C)** The same data are plotted, as in panel A with the addition of the average single unit SDF for phase locked PFC single units (dark blue line).

Having demonstrated that both PFC spiking and PFC hGamma power co-fluctuate around the peak of LC-MUA 5 Hz oscillations, we predicted that PFC single units would be phase-locked to LC-MUA 5 Hz oscillations. In four of the rats shown in **Figure 3**, we used a 4-shank silicone probe (200 um between shanks) placed in the anterior-posterior plane within the PFC deep layers. These probes were chosen to isolate PFC single units. Each shank had 2 recording tetrodes separated by 500 um in the dorsal-ventral direction. Using these probes, we recorded 83 PFC single units (S.E.M.: 17±2 units per rat, Range: 9 to 22 units). Single unit spike trains were converted to SDFs using a 100 msec Gaussian kernel. In line with this prediction, we found that 20 of 83 PFC single units (27%) were significantly phase-locked to LC-MUA 5 Hz oscillations (Rayleigh’s test for circular uniformity, p<0.05). The phase preference of these PFC units concentrated around the trough of the LC-MUA 5 Hz oscillation (**Figure 6B**). The spike rate of the PFC single units, which were phase locked to LC-MUA 5Hz oscillation, increased ~100 msec prior the LC-MUA peak (**Figure 6C**). The PFC spike rate increased ~100 msec prior to the LC-MUA peak (purple line, **Figure 6C**). This delay is inconsistent with the known conduction delays (~29 msec) from the PFC to the LC (Jodo, et al. 1998). Our findings suggest that the spiking of some PFC single units has a temporally consistent relationship with the LC-MUA 5 Hz oscillation, but that these spikes occur far earlier (~100 msec) in the LC-MUA oscillation cycle to monosynaptically drive its ascending phase given conduction delay of ~29 msec. Although poly-synaptic influence of these PFC spikes on LC could not be ruled out, our sample of PFC units does not support the claim that PFC spike output monosynaptically drives the 5 Hz rhythmic firing in LC.

The role of PFC spikes phase-locked to the trough of the LC-MUA 5 Hz oscillation remains unclear (**Figure 6B**). The firing of a subset of PFC single units during the trough of the LC-MUA 5 Hz oscillation suggest that they may have a role in the rhythmic prefrontal-coeruleus interaction. One possibility is that an increase in PFC spiking ~100 msec prior to the LC-MUA peak is a local circuit mechanism that drives both the hGamma power increase and synchronous PFC spiking. We examined this possibility by calculating TE between PFC hGamma amplitude and PFC single units phase locked to LC-MUA 5 Hz oscillation (**Figure 7**). We found that PFC spiking was predictive of the upcoming hGamma power increase (Wilcoxon rank sum test due to lack of normal distribution, Z = 2.61, p = 0.009, Cohen’s D = 0.776, Power = 0.736). The spiking of PFC units with no consistent relation to LC-MUA 5 Hz oscillation (Rayleigh’s test for circular uniformity, p>0.2) was not predictive of the hGamma power change (Wilcoxon rank sum test, Z = 1.429, p = 0.153, Cohen’s D = 0.343, Power = 0.336).

**Figure 7.**
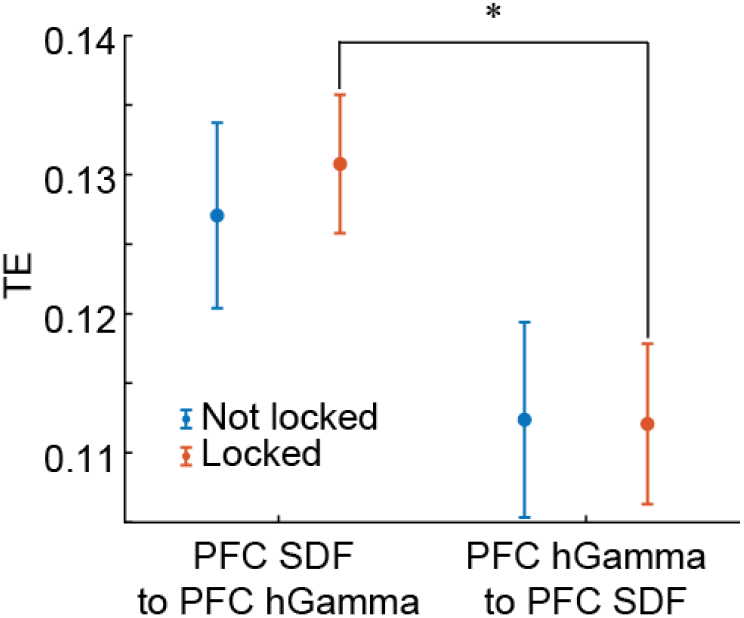
The spiking of PFC single units that are phase locked to LC-MUA 5 Hz oscillation predict the local PFC hGamma power increase. Transfer entropy (TE) between the PFC hGamma amplitude and PFC SDF was higher in the direction of the spiking to the hGamma signal. This directionality difference in TE was significant only for the units that were phase locked to LC activity.

### The PFC unit-pair spike synchrony during LC-PFC interactions

Synchronous spiking between PFC neurons could be an alternative mechanism mediating the PFC effects on the LC. We tested the possibility that the increases in PFC hGamma power are associated with a transient increase in synchronous spiking across PFC single units. We constructed joint peri-event time histograms of PFC single unit spiking using a ±60 msec window around PFC hGamma transients (Aertsen et al., 1989; Brody, 1999). The joint peri-event spike histogram was calculated in 10 msec bins in order to capture spiking synchronized across single units with enough temporal proximity to evoke post-synaptic effects on target neurons (Abeles, 1982; Alonso et al., 1996; Fujisawa et al., 2008). The diagonal of the joint peri-event time histogram was used to calculate a coincidence histogram for each of the 808 pairs of PFC single units. The coincidence histograms serve to characterize synchronous spiking around the time of the hGamma transient (t = 0 in **Figure 8**). We found an increase in synchrony around PFC hGamma power peaks in half (48%) of PFC single unit pairs (**Figure 8**). The synchronous spiking occurred ~ 20 msec prior to the hGamma power peak and lasted for ~ 50 msec. The hGamma power peak itself occurred ~ 29 msec prior to the LC-MUA 5 Hz peak, which means that synchronous PFC spiking transiently increased ~ 49 msec before the LC-MUA 5 Hz peak and lasted ~ 21 msec after the LC-MUA 5 Hz peak. Note that PFC spikes occurring 20 msec after the hGamma peak can still drive the LC-MUA during the descending phase of the LC-MUA 5Hz oscillation given prior work demonstrating the PFC-to-LC monosynaptic conduction time as fast as 10 msec for some units (Jodo, et al. 1989). These data suggest a potential mechanism for PFC monosynaptic control over LC firing using transiently synchronous spiking during hGamma transients. However, it is important to note that the recorded PFC single units may or may not project to the LC. In summary, it appears that the subset of PFC neurons that spike during the troughs of LC-MUA 5 Hz oscillations may drive local hGamma transients, which are themselves related to the synchronous spiking of PFC units that may drive the LC-MUA increase.

**Figure 8.**
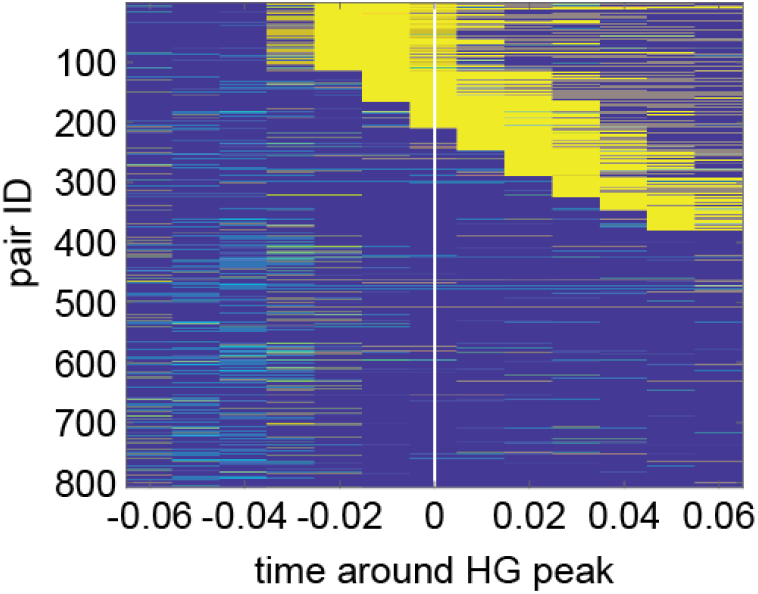
PFC single unit-pair synchrony increases during PFC hGamma transients. The coincident histograms of 808 PFC single unit pairs (y-axis, sorted by synchrony onset time) show an increase in pairwise unit spike synchrony around PFC hGamma power peaks (x-axis). The coincident histograms are z-scored with increases in synchrony (Z-score greater than 2) in yellow.

## Discussion

In this study, we examined the relationship between rhythmic (~5 Hz) increases in LC-MUA and neural activity in the PFC, an important forebrain target of LC. In contrast to the well-described slower (~1 - 2 Hz) rhythmic increases in LC spiking that are observed during cortical slow oscillations (Eschenko et al., 2012; Safaai et al., 2015; Totah et al., 2018), the LC-MUA 5 Hz oscillations predominantly occurred during the activated cortical state devoid of cortical slow oscillations. By measuring cross-frequency coupling between LC-MUA oscillations within 1 - 10 Hz range and the power spectrum of PFC LFP (30 – 500 Hz), we observed a systematic temporal relationship between the phase of LC-MUA oscillations within a 4 - 6 Hz range and PFC hGamma power (60 - 200 Hz). The transient increase in PFC hGamma power preceded the LC-MUA 5 Hz oscillation peak by ~29 msec. This time lag is consistent with the previously reported orthodromic conduction times from deep layers of the prelimbic division of the PFC in rats (Jodo et al., 1998). Furthermore, we provided evidence for causal influence of PFC hGamma on LC-MUA during episodes when LC population firing rate rhythmically fluctuated at ~5 Hz. Namely, we observed higher transfer entropy from PFC hGamma power to LC-MUA 5 Hz phase than in the opposing direction. Together, these findings suggest that PFC hGamma transients may predict excitability of LC.

hGamma transients are unlikely to have a direct synaptic effect on LC neurons because they are highly local. We showed that synchronous spiking between PFC single units occurs during hGamma transients and reached maximum around the peak of LC-MUA 5 Hz oscillation. We suggest that this increased population synchrony in PFC may be top-down excitatory input to LC. The timing between synchronous PFC spiking and the peak of the LC-MUA oscillation is consistent with the conduction velocity of the prefrontal-coeruleus projection (Jodo et al., 1989). Synchronous spiking is an ideal candidate for top-down glutamatergic control over LC neurons because glutamatergic neurons spiking within ~5 msec of one another evoke a larger post-synaptic response (Abeles, 1982; Alonso et al., 1996; Fujisawa et al., 2008). Collectively, our findings suggest that the PFC hGamma transients and, critically, associated neuronal spike synchrony may be a sign of PFC top-down control over LC population activity.

We also observed a subpopulation of PFC single units (~27%) that increased their firing rate ~100 msec prior to the peak of LC-MUA 5 Hz oscillation. Given the conduction velocity of the prefrontal-coeruleus projection, these spikes are unlikely to monosynaptically drive LC-MUA. However, their consistent timing in relation to the trough of LC-MUA 5 Hz oscillation suggested that these spikes are involved in the prefrontal-coerulear interaction. Transfer entropy analysis revealed that the spikes of these PFC neurons were predictive of the upcoming change in PFC hGamma power. Therefore, a sub-set of PFC single units which are phase locked to the trough of the LC-MUA 5 Hz oscillation (thus preceding the hGamma power increase) may drive the hGamma transient and the associated synchronized spiking in PFC. It also cannot be excluded that PFC spikes consistently occurring ~100 msec prior to the LC-MUA peak could drive LC-MUA directly via multiple polysynaptic routes.

Overall, we propose that a local PFC circuit mechanism drives the synchronous spiking that influences LC-MUA (**Figure 9**). It remains to be determined how the 5 Hz rhythmicity in the LC emerges and if it is specific to anesthesia or has functional significance in behaving animals.

**Figure 9.**
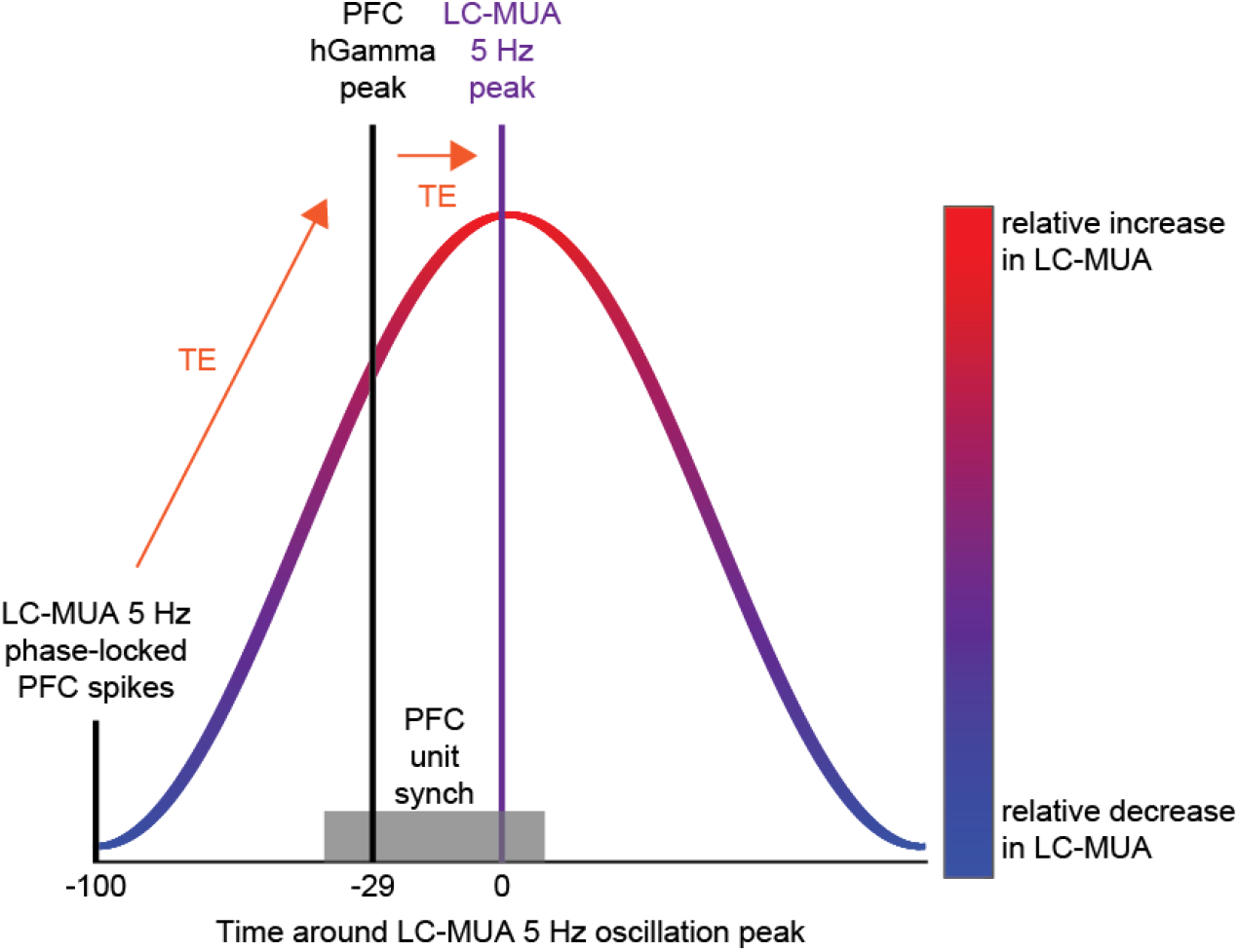
A putative model of top-down prefrontal control over the LC. We found that PFC spikes were phase locked to the trough of LC-MUA 5 Hz oscillation. Given the 10 to 70 msec conduction time from the PFC to the LC (Jodo, et al. 1989), this time point is too early to conduct a signal that could evoke an increase in LC-MUA spike rate (red part of the oscillation). Instead, these PFC spikes were predictive of a local hGamma power increase (orange arrow, direction of transfer entropy, TE). This hGamma transient was, in turn, predictive of the subsequently increased LC-MUA (red peak). Data are shown in **Figure 4A**. The hGamma transient precedes the LC-MUA peak by 29 msec. Data are shown in **Figure 3C** and **Figure 4B**. In a window of −20 msec to + 50 msec around the hGamma peak (or −49 msec to +21 msec around the LC-MUA peak), PFC single unit pairs spike with transiently increased synchrony (grey area on x-axis). Data are shown in **Figure 8**. A chain of neural events from the PFC spikes time locked to the through of LC-MUA 5 Hz oscillation to hGamma-associated spike synchrony in the PFC may drive an increase in LC spike rate.

## Methods

### Experimental Model And Subject Details

#### Subjects

All experimental procedures were carried out with approval from the local authorities and in compliance with the German Law for the Protection of Animals in experimental research (Tierschutzversuchstierverordnung) and the European Community Guidelines for the Care and Use of Laboratory Animals (EU Directive 2010/63/EU). Male Sprague-Dawley rats (350 - 450 g) were used. Animals (specific pathogen free) were ordered from Charles River Laboratories (Sulzfeld, Germany). Animals were pair housed and on a 08:00 to 20:00 dark to light cycle. Data were collected from rats used in a prior study (Totah et al., 2018).

### Method Details

#### Anesthesia and Surgical Procedures

Rats were anesthetized using an intra-peritoneal (i.p.) injection of urethane at a dose of 1.5 g/kg body weight (Sigma-Aldrich, U2500). Oxygen was administered throughout the procedure and body temperature was maintained at 37 C using a heating pad and rectal probe to monitor body temperature. The skull was leveled to 0 degrees, such that the difference between lambda and bregma was less than 0.2 mm.

#### Stereotaxic coordinates and electrode placement

Electrodes were targeted for the LC and PL. The coordinates for LC were 4.0 mm posterior from lambda, 1.2 mm lateral from lambda, and approximately 6.0 mm ventral from the brain surface (implanted at a 15 deg posterior angle). The following coordinates, in relation to bregma and the brain surface, were used for PL: 3.0 mm anterior, 0.8 mm lateral, 3.0 mm ventral.

The LC electrode was targeted based on standard electrophysiological criteria. These criteria included a slow spontaneous firing rate, biphasic response to noxious sensory stimuli (foot shock), audible presence of jaw movement-responsive cells in the Mesencephalic Nucleus of Cranial Nerve V with undetectable single units (<0.2 mV) from that structure. LC electrode placements were also verified using histological examination in 50 um sections that were stained for Cresyl violet or a DAB and horse radish peroxidase reaction with hydrogen peroxide to visualize an antibody against tyrosine hydroxylase (the catecholamine synthesis enzyme).

#### Electrodes

The LC was recorded either using a single tungsten probe (FHC, Model: UEWMFGSMCNNG) or a multi-channel silicone probe (NeuroNexus, Model: A1×32-Poly3-10mm-25s-177-A32). Deep layer PFC LFP was recorded using a single tungsten probe (FHC). The impedance was 200 kOhm to 800 kOhm. For recordings of PFC single units, a Neuronexus A4×2-tet-5mm-500-400-312 probe was used. The probe was oriented running anterior-posterior in the deep layers.

#### Recording and signal acquisition

A silver wire inserted into the neck muscle was used as a reference for the electrodes. Electrodes were connected to a pre-amplifier (in-house constructed) via low noise cables. Analog signals were amplified (by 2000 for LC and 500 for cortex) and filtered (8 kHz low pass, DC high pass) using an Alpha-Omega multi-channel processor (Alpha-Omega, Model: MPC Plus). Signals were then digitized at 24 kHz using a data acquisition device CED, Model: Power1401mkII).

#### Administration of clonidine

At the end of the recording, a 0.05 mg/kg dose of the alpha-2 adrenergic agonist clonidine was injected i.p. (Sigma-Aldrich, product identification: C7897). The recording was continued at least until LC activity ceased.

#### Determination of cortical state

Cortical states were separated based on characteristics of the LFP signal examined in 7 sec windows. Two characteristics were considered: a ratio of the cortical LFP power below 4 Hz and the power above 20 Hz and the kurtosis of the distribution of LFP values. The LFP was first decimated and low-pass filtered to 500 Hz. The distribution of power ratio values and kurtosis values for each 7 sec window were fit with Gaussian mixture models. We used the power ratio to label windows of data as putative activated states if they were <1 standard deviation from the lower Gaussian’s mean or they were labeled putative slow oscillation states if they were >-1 standard deviation from the higher Gaussian’s mean. We used the kurtosis values to label windows of data as putative activated states if they were >1 standard deviation from the higher Gaussian’s mean or as putative slow oscillation states if they were < 1 standard deviation from the lower Gaussian’s mean. Any labels that agreed across the kurtosis-based labels and the power ratio-based labels were used as the final state assignments for those windows. Any windows that were unlabeled or did not agree across the two characteristics were ignored in order to conservatively reduce mistaken classifications. The raw LFP signals were plotted for visual inspection in order to assess the accuracy of labeling.

#### Detection of LC MUA oscillations

The LFP (digitized and stored at 24 kHz) recorded in the LC was bandpass filtered for high frequency, spiking activity (400 to 3000 Hz) to obtain a multi-unit spiking signal, as would be done typically for sorting single unit spikes. The signal was downsampled to 9 kHz. The signal was then rectified. This signal is termed the MUA signal. The power spectral density (PSD) of the MUA signal was obtained using a multi-taper estimation generated from the Chronux toolbox in MATLAB (params.tapers = [3 5]). For each recording session (one per rat), the PSD of LC-MUA was calculated in 4 sec windows. The resulting PSD’s were k-means clustered. An optimal k was determined by a gap statistic. The mean PSD for each cluster was plotted and manually inspected for a 4 – 7 Hz peak. In some cases, multiple clusters of PSDs had a peak in the 4 – 7 Hz range with the only difference being the amplitude of the spectral peak. For each recording session, all clusters with 4 - 7 Hz peak were accepted as epochs with LC-MUA 5 Hz oscillations.

#### Cortical spectrogram calculation

Cortical spectrograms, triggered on LC MUA oscillation peaks, were calculated as follows. The LC MUA signal was bandpass filtered 5 Hz peak frequency (4 – 6 Hz) and Hilbert transformed to obtain the instantaneous phase. We selected peak times that occurred during the 4 sec windows with 5 Hz oscillations (defined by PSD clustering, see above section). A cortical spectrogram was generated for ±5 sec around this peak using a complex Morelet wavelet transform. The large window was used to discount edge artifacts. The resulting analytic amplitudes were then cut to a small time around the oscillation. At each cortical frequency, the spectrogram was normalized as a Z-score. The normalization was done around each LC MUA oscillation peak, then averaged across peaks for each rat. The presented spectrograms are the averages across rats.

#### Coupling between LC MUA oscillation phase and cortical LFP oscillation amplitude

The phase-amplitude coupling was calculated using the LC MUA signal as the oscillation for phase and the cortical LFP signal (downsampled to 9 kHz) as the oscillation for amplitude. The relationship between phase of one frequency and the amplitude of another frequency was quantified using the modulation index (MI), which is based on the Kullback-Leibler divergence of the circular distribution from uniformity (Tort et al., 2010). MI was calculated for each frequency pair (a frequency for phase, *f_P_*, and a frequency for amplitude, *f_A_*). Only *f_A_* that were two times *f_P_* were considered, so that phase of at least two oscillation cycles was present for measuring the MI. We binned phase into 18 bins, where *j* is a bin, and then calculated the mean amplitude, 〈*A_f_A__〉*) of 〈*A_f_A__*〉_*θ*_*f_P_*_(*i*)_. We normalized the distribution by dividing each bin by the sum across all bins. The resulting distribution is

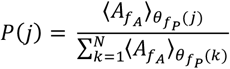

where *k* is the phase bin and *N* is the total number of phase bins. The third step was to quantify the difference of this amplitude distribution from a uniform circular distribution. This was done using the Kullback-Leibler divergence. The first step in calculating the divergence was to calculate the Shannon Entropy of *P*(*j*), which is

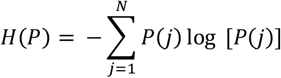

The second step was to calculate the Kullback-Leibler divergence of the amplitude distribution from a uniform distribution, which is related to Shannon Entropy as,

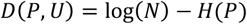

where *U* is the uniform circular distribution. Note that, if the amplitude distribution is flat and the amplitude of *f_A_* is the same for all phase bins of *f_P_*, then log (*N*) is the maximal possible entropy in which *P*(*j*) = 1/*N* and phase is equally distributed across all bins, *j*. Accordingly, the Kullback-Leibler divergence is normalized by the maximal entropy, log (*N*), in which case a uniformly distributed *P*(*j*) that is not different from *U* will push the MI to 0. Otherwise, MI will range 0 to 1, with 1 indicating that oscillations of *f_A_* exist in a single *f_P_*(*j*). The MI is thus,

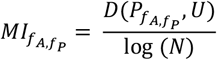

In order to control for chance modulation, we constructed a surrogate set of MI values to measure the level of coupling between *f_A_* and *f_P_* that could occur by chance. We shuffled *f_A_* and then calculated a surrogate MI. We performed this procedure 100 times. A 99% confidence interval threshold was subtracted from the MI of the real data, such that values equal to or less than 0 were non-significant.

#### PFC single unit spike sorting

Single unit spike sorting was performed using MountainSort (Chung et al., 2017). Units were assessed for amplitude stability over time, a low proportion (<one quarter of the shoulder of the auto-correlogram) of spikes in the ±1 msec interval of the auto-correlogram, and cross-correlograms not indicative of recordings from the same unit split into multiple clusters.

#### Joint peri-event time histograms

The joint peri-event time histograms were normalized by subtracting the top 5% value obtained by selecting random event times that were equivalent to the number of hGamma events. We plotted the coincidence histogram using the values along the ±30 msec diagonal of the joint peri-event time histogram. These values were chosen because the hGamma transients to which the histograms were aligned lasted about 60 msec.

## Data And Software Availability

The generated datasets and MATLAB codes for analysis are available from the Lead Contact on reasonable request.

## Acknowledgements

We thank Prof. Stefano Panzeri and Dr. Michel Besserve for help implementing the transfer entropy analysis in MATLAB. We are grateful to Dr. Antonio Fernández-Ruiz and Dr. Martin Vinck for comments on the manuscript.

## Notes

### Competing Interest Statement

The authors have declared no competing interest.

